# PERK-mediated induction of miR-5p and miR-3p arms of miR-616 regulates cell growth by targeting c-MYC

**DOI:** 10.1101/2023.03.06.531445

**Authors:** Vahid Arabkari, Afrin Sultana, David Barua, Mark Webber, Terry Smith, Ananya Gupta, Sanjeev Gupta

## Abstract

C/EBP homologous protein (CHOP), also known as DNA damage-inducible transcript 3 (DDIT3), is a member of CCAAT/enhancer-binding protein (C/EBP) family. The expression of CHOP is upregulated during unfolded protein response (UPR), and sustained CHOP activity plays an important role in UPR-induced apoptosis. MicroRNA-616 is localized in an intron of the CHOP gene. However, regulation of miR-616 expression during UPR and its function in breast cancer is not clearly understood. We show that miR-5p/-3p arms of miR-616 are expressed with levels of 5p arm higher than 3p arm. During conditions of UPR, the expression of miR-5p and miR-3p arms of miR-616 and its host gene (CHOP) was concordantly increased primarily in a PERK-dependent manner. We show that ectopic expression of miR-616 significantly suppressed cell growth and colony formation, whereas knockout of miR-616 increased it. We identified that MYC proto-oncogene (c-MYC) gene is repressed during the UPR and targeted by miR-616. Further, we show that expression of miR-616 and CHOP is downregulated in human breast cancer, where expression of miR-616 was associated with poor overall survival (OS) in luminal A subtype and better OS HER2 subtype of breast cancer. In summary, our results suggest a dual function for the DDIT3 locus, where CHOP protein and miR-616 can co-operate to regulate cancer progression.

## INTRODUCTION

The endoplasmic reticulum (ER) is a crucial cellular organelle that plays important role in folding and maturation of proteins that transit through the secretory pathway. Stimuli that compromise the protein handling capacity of the ER activate an evolutionarily conserved pathway known as unfolded protein response (UPR). Three major sensor molecules that initiate the UPR are activating transcription factor 6 (ATF6), inositol-requiring enzyme 1α (IRE1) and protein kinase RNA-like endoplasmic reticulum kinase (PERK) (1). During resting state the activity of PERK, IRE1 and ATF6 is kept under control by interaction with ER chaperone glucose-regulated protein 78 (GRP78); however, upon accumulation of misfolded proteins in ER, GRP78 dissociates from these molecules, leading to their activation (2). Upon dissociation with GRP78, IRE1 auto-phosphorylates and oligomerizes leading to the activation of its endoribonuclease domain present in the cytosolic domain. Active IRE1 catalyses the unconventional splicing of XBP1 mRNA and regulated IRE1-dependent decay of transcripts (RIDD) (3). While splicing of XBP1 mRNA is an adaptive response to UPR, RIDD has many context dependent outcomes (3). ATF6 is a transmembrane protein located in the ER membrane with an N-terminus cytoplasmic domain containing a DNA-binding motif and a C-terminus ER-luminal domain that interacts with GRP78 (4). During UPR, ATF6 is relocates from the ER to the Golgi compartment. In the Golgi, ATF6 is processed by intramembranous proteolysis causing the release of the transcriptionally active N-terminal cytosolic domain, p50ATF6 (5). During UPR, active PERK phosphorylates eukaryotic initiation factor 2α (eIF2α), causing reduced translation initiation for a majority of cellular proteins and preferential translation of a subset of mRNAs including activating transcription factor 4 (ATF4) (6, 7). Phosphorylation of eIF2α leads to general translational block and preferential translation of a subset of mRNAs including activating transcription factor 4 (ATF4) (7). Phosphorylation of NRF2 by PERK attenuates Keap1-mediated degradation of NRF2 and promotes expression of NRF2-target genes through the antioxidant response elements (ARE) (6). The increased protein handling capacity of the ER and degradation of unfolded proteins induced by UPR attempts to restore the protein homeostasis and increase cell survival. However, if UPR-induced mechanisms fail to alleviate ER stress, UPR activates apoptosis.

MicroRNAs (miRNAs) are small (∼20–23 nucleotide) single-stranded RNA molecules that regulate gene expression in a sequence-specific manner (8). The increased expression of miR-616 has been reported in androgen-independent prostate cancer (9, 10), gastric cancer (11), non-small cell lung cancer (NSCLC) (12), hepatocellular carcinoma (HCC) (13) and gliomas (14). MiR-616 targets tissue factor pathway inhibitor (TFPI-2) in prostate cancer (9); PTEN in HCC and gastric cancer (15); SOX7 in glioma and NSCLC (14). However, miR-616-3p has been reported to reduce XIAP expression and potentiate apoptosis in HUVECs (16). LINC01614 promotes head and neck squamous cell carcinoma progression via PI3K/AKT signalling pathway and miR-616-3p can inhibit cancer progression by downregulating LINC01614 expression (17). Ye et al. reported that miR-616-3p inhibited cell growth and mammosphere formation of breast cancer cells by suppressing GLI1 (18), while Yuan reported that miR-616-3p promoted breast cancer cell migration and invasion by targeting TIMP2 and regulating MMP signalling (19). Overall, these observations suggest that miR-616 exhibits both oncogenic and tumour suppressor functions in context- and/or tissue-dependent manner. Additional studies are warranted to further investigate the role of miR-616 in human cancers.

During the past few years, work from several groups has revealed that all three branches of the UPR regulate specific subsets of miRNAs (20, 21, 22, 23). The modes of regulation include both increase and decrease in expression by UPR-regulated transcription factors such as ATF6, XBP1, ATF4 and NRF2 as well as degradation by RIDD activity. The key feature of UPR-dependent miRNA expression is fine tuning of the ER homeostasis to modulate cellular adaptation to stress and regulation of cell fate. The gene encoding miRNA-616 is located in the second intron of CHOP/DDIT3. This genomic co-location of miR-616 with CHOP/DDIT3 suggests a role for miR-616 in UPR. Indeed, increased expression of miR-616 has been reported during conditions of UPR (24, 25). Hiramatsu et al., focused on the expression of miR-616-5p (24) while Arabkari focused on the expression of miR-616-3p (25). Here we have evaluated the expression of both miR-5p/-3p arms of miR-616 as well its host gene (CHOP) during conditions of UPR.

We show that expression level of miR-616-5p was higher than miR-616-3p in human cancer cells and tissues. During conditions of UPR, the expression of both miR-5p/-3p arms of miR-616 and CHOP was increased in a PERK-dependent manner. Our results show that miR-616 regulates cell growth and proliferation along with regulation of c-MYC expression. We identified that MYC proto-oncogene (MYC) gene is repressed during the UPR and targeted by miR-616. Finally, we show that expression of miR-616 and CHOP (host gene of miR-616) is reduced in human breast cancer. In summary, our results suggest that CHOP locus generates two gene products, where CHOP protein and miR-616 can act together to regulate cancer progression.

## METHODS

### Cell culture and treatments

Human breast cancer cells (MCF7 and BT474) were purchased from ECACC. HCT116 cells were kind gift from Dr. Victor E. Velculescu, Johns Hopkins University, USA. HEK 293T cells were gift from Indiana University National Gene Vector Biorepository. The MCF7, BT474 and HEK 293T cells were maintained in Dulbecco’s modified eagle’s medium. HCT116 cells were maintained in McCoy’s 5A modified. The medium was supplemented with 10% heat inactivated foetal bovine serum (FBS) and 100 U/ml penicillin and 100 μg/ml streptomycin with 5% CO2 at 37°C. Thapsigargin (Cat # 1138) and BFA (Cat # 1231) were purchased from Tocris Bioscience. All reagents were purchased from Sigma-Aldrich unless otherwise stated.

### Plasmid constructs

miExpressTM precursor miRNA expression clones, miR-CTRL and miR-616 (pEZX-MR03 vector) were sourced from GeneCopoeia, Rockville, MD, USA. The lentiviral plasmids pLV [CRISPR]-hCas9:T2A: Puro-U6>mir-616 expressing hCas9 protein and miR-616 targeting gRNA were obtained from VectorBuilder Inc, Chicago, IL, USA. The sequences of two miR-616 targeting gRNAs are 5’-GGAAATAGGAAGTCATTGGA-3’ and 5’-GTGTCATGGAAGTCACTGAA-3’. PCDH-Flag-c-MYC was a gift from Hening Lin (Addgene plasmid # 102626; http://n2t.net/addgene:102626; RRID: Addgene_102626).

### Generation of stable cell lines

MCF7 cells were transduced with lentiviruses to generate stable miR-616 overexpressing and miR-616 knockout sub-clones. Lentivirus was generated by transfecting lentiviral plasmids along with packaging plasmids in 293T cells using jetPEI transfection reagent as described previously (26).

### RNA extraction, Reverse transcription reaction and real time quantitative PCR

Total RNA was isolated using Trizol (Life Technologies) according to the manufacturer’s instructions. RPLP0 and GAPDH (for mRNAs) and U6 SnRNA (for miRNAs) were used as a reference genes to determine the relative expression level of target genes between treated and control samples using△ △Ct method.

### Colony formation assay

The cells were plated in 6 well plates and were grown for 14 days. After 14 days, cells were washed twice with PBS and fixed with 10% formaldehyde for 5 minutes and stained with 0.5 % crystal violet for 10 minutes. The number of colonies were counted in five random view fields under a microscope and average number of colonies was determined. Colony size was determined using Image J software.

### X-CELLigence cell proliferation assay

xCELLigence experiments were performed using the Real-Time Cell Analyzer (RTCA) Dual Plate (DP) instrument according to the manufacturer’s instructions (Agilent.com). Briefly, instrument-specific specially designed gold microelectrodes fused microtiter 16-well cell culture plate (0.2 cm2 well surface area; 250 μl volume per well) was used to monitor the real-time changes expressed as Cell index (CI). Cell Index is defined as (Rn−Rb)/15 where Rn is the cell–electrode impedance of the well with the cells and Rb is the background impedance of the well with the medium alone. The background impedance was measured by adding 50 μl of medium to each well of the E-Plate before seeding the cells. After seeding (2500 cells/well), the E-plate was incubated at room temperature for 30 min and then transferred to the instrument. Cell proliferation was monitored every 15 min for 50 h.

### Western blot analysis

Western blotting procedure has been described previously. After blocking with specific solution nitrocellulose membranes were treated with specific primary antibodies including c-MYC (Santa Cruz Biotechnology Cat# sc-40), ERα (Merck Millipore Cat# 2828587) and β-Actin (Sigma, Cat# A-5060) at 4°C for overnight. After washing three times with PBS/0.05%Tween solution, the membranes were incubated with appropriate secondary antibody at room temperature for 2 hours. The membranes were then washed twice with PBS/0.05%Tween and once with PBS and finally the signals were detected using Western Lightening chemiluminescent substrate (Perkin Elmer, Netherlands, Cat #NEL104001EA).

### Statistical Analysis

The GraphPad was used for statistical analysis and generating graphs. Two-tailed unpaired t-test was performed to determine statistically significant differences between independent groups. Results with a p<0.05 were considered statistically significant.

## Results

### Expression of miR-616-5p is higher than miR-616-3p

Several miRNAs are located within the introns of either protein coding or non-coding genes, and are referred to as intronic miRNAs (27). The genes which harbour these miRNAs are embedded are called host genes. MiR-616 is an intronic miRNA localized in the first intron of DDIT3 gene (**Fig 1A**). Both the 5’ and 3’ arms of the precursor duplex can generate mature miRNAs, and are referred to as miRNA-5p and -3p (27, 28). The miRBase (http://www.mirbase.org) (29) has discontinued the use of miRNA/miRNA* nomenclature and use of miR-5p and miR-3p nomenclature (30), based solely on 5’- or 3’-arm derivation of the mature miRNA is now recommended. The sequences of mature miR-616, accession numbers and the previous names of miR-616-5p and -3p as per miRNA/miRNA* nomenclature are shown in Fig 1B. Indeed, both 5p and 3p arms of miR-616 have been shown to be expressed in human cancers (11, 31). Next we evaluated the expression of miR-616-5p and miR-616-3p in human cancers. Initially expression of miR-616-5p and miR-616-3p was determined in various human cancer cell lines by qRT-PCR. We observed that endogenous expression of miR-616-5p was higher than miR-616-3p in all the cell lines tested (**Fig 1C**). Next we analysed the expression of miR-616-5p and miR-616-3p in the TCGA dataset from different human cancers using OncomiR (http://www.oncomir.org), a web-based tool for miRNA expression analysis. We observed a higher expression of miR-616-5p as compared to miR-616-3p in all the cancer types evaluated (**Fig 2**). Of note the ratio of miR-616-5p and miR-616-3p expression varied considerably across the different tissue types. These results suggest that both 5p and 3p arms of miR-616 are expressed where miR-616-5p accumulates at higher levels than miR-616-3p but their ratio is controlled in tissue-specific manner.

**Fig 1.**
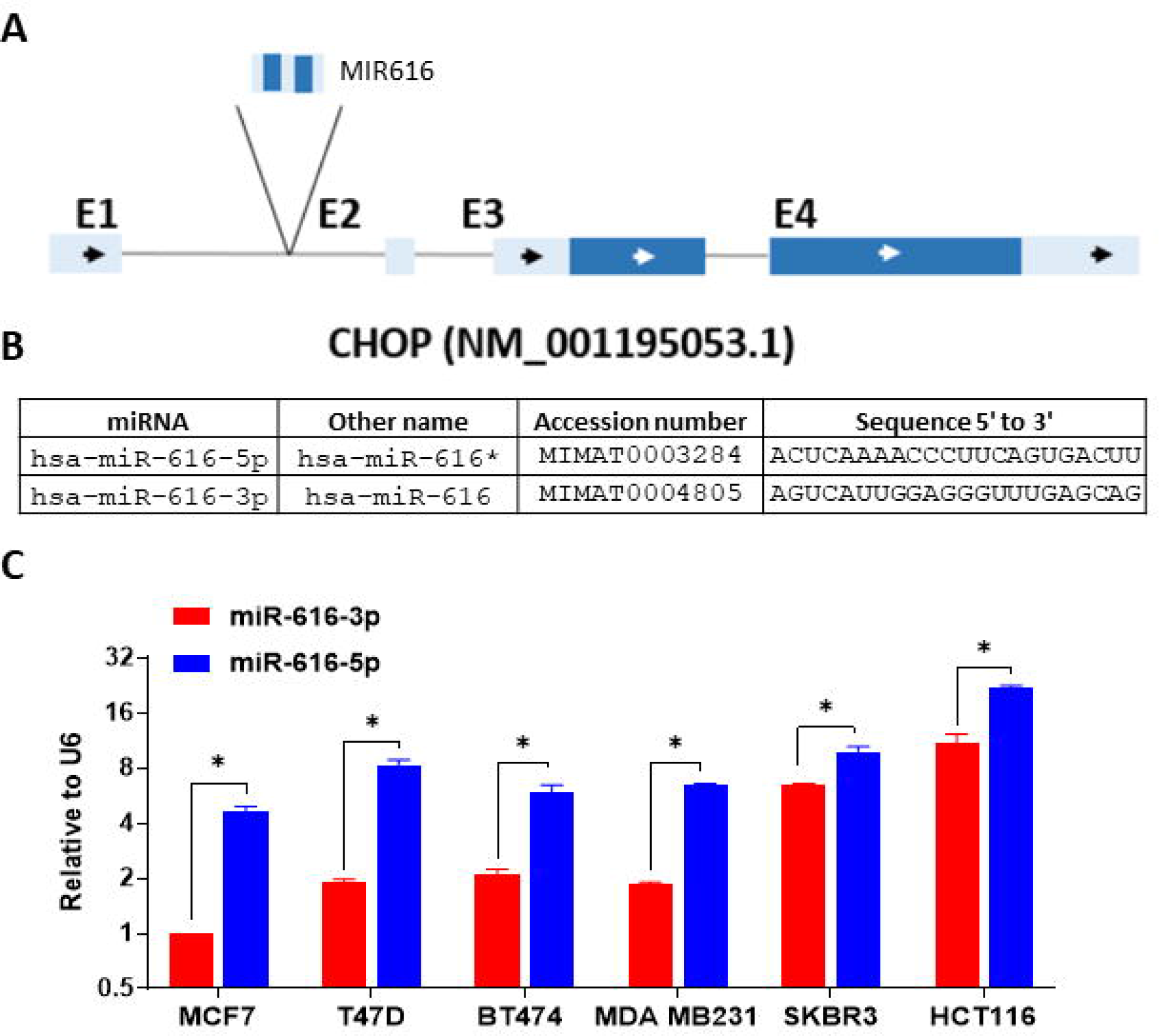
Basal expression of miR-616-5p and miR-616-3p in human cancer cell lines. (A) Schematic representation of the CHOP locus is shown. The exons are shown in blue and gray line between them represents intron. The miR-616 is located in the first intron. The arrows show the direction of transcription of the gene. The protein coding regions are shown in dark blue and untranslated regions are shown in light blue. (B) The name, accession number and sequence of miR-616-5p and 3p are shown. (C) Total RNA from indicated cells was used to determine the expression of miR-616-5p and miR-616-3p by qRT-PCR and normalised against RNU6. Expression of miR-616-3p in MCF7 cells was arbitrarily set at 1. Error bars represent mean ±S.D. from three independent experiments performed in triplicate. *P < 0.05, two-tailed unpaired t-test compared to miR-616-3p in MCF7 cells.

**Fig 2.**
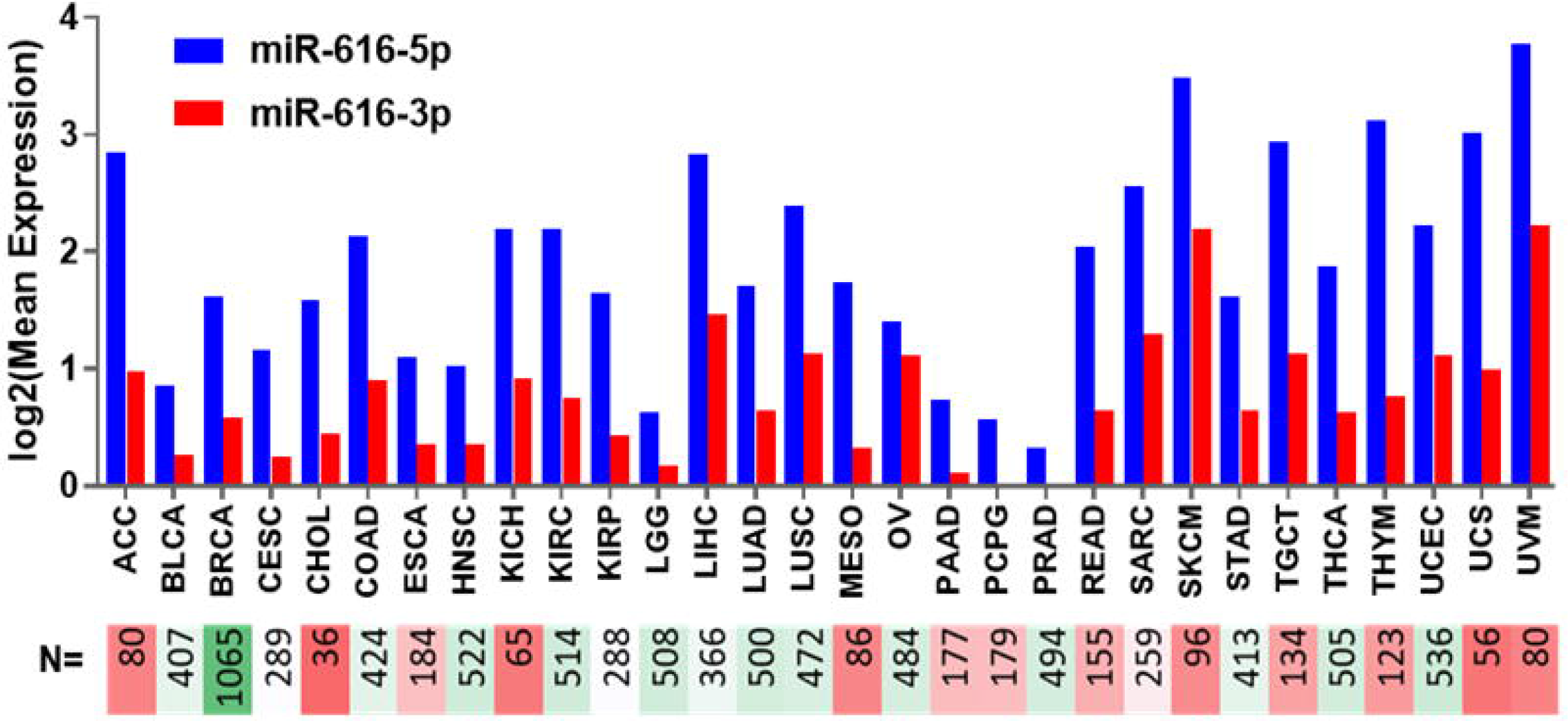
Expression of miR-616-5p and miR-616-3p in human cancers. Web-based algorithm at (http://www.oncomir.org/) was used to assess the expression of miR-616-5p and miR-616-3p. Log2 mean expression of miR-616-5p and 3p in indicated cancer types is shown. (N = number samples in each cancer type). ACC: adrenocortical carcinoma; BLCA: bladder urothelial carcinoma; BRCA: breast invasive carcinoma; CESC: cervical squamous cell carcinoma and endocervical adenocarcinoma; CHOL: cholangiocarcinoma; COAD: colon adenocarcinoma; ESCA: esophageal carcinoma; HNSC: head and neck squamous cell carcinoma; KICH: kidney chromophobe; KIRC: kidney renal clear cell carcinoma; KIRP: kidney renal papillary cell carcinoma; LGG: brain lower grade glioma; LIHC: liver hepatocellular carcinoma; LUAD: lung adenocarcinoma; LUSC: lung squamous cell carcinoma; MESO: mesothelioma; OV: ovarian serous cystadenocarcinoma; PAAD: pancreatic adenocarcinoma; PCPG: pheochromocytoma and paraganglioma; PRAD: prostate adenocarcinoma; READ: rectal adenocarcinoma; SARC: sarcoma; SKCM: skin cutaneous melanoma; STAD: stomach adenocarcinoma; TGCT: testicular germ cell tumours; THCA: thyroid carcinoma; THYM: thymoma; UCEC: uterine corpus endometrial carcinoma; UCS: uterine carcinosarcoma; UVM: uveal melanoma

### Increased expression of miR-616-5p and miR-616-3p during UPR

It was initially proposed that intronic miRNAs are derived from the same primary transcript as their host genes thereby leading to co-expression and co-regulation of miRNA and cognate host gene (27). Analysis of expression of 175 miRNAs and their host genes across twenty four different human organs reported a significantly correlated expression profile (32). Recent evidence however, shows significant discordant expression between intronic miRNAs and host genes. Two likely explanations for this discordant expression are: (i) miRNAs have their own independent promoters and (ii) crosstalk between microprocessor cleavage and mRNA splicing (33, 34). Next, we evaluated the co-regulation of miR-616 and its host gene (CHOP) during conditions of UPR. MCF7, BT474 and HCT116 cells were either treated or untreated with thapsigargin (TG) or Brefeldin A (BFA) for 24 hours and expression of CHOP, miR-616-5p and miR-616-3p was determined. The expression level of CHOP, miR-616-5p and miR-616-3p was upregulated in all three cell lines tested, during ER stress (**Fig. 3A-C**). Next we investigated the role of ATF6, PERK and IRE1, three key mediators of the UPR in upregulation of miR-616 expression. For this purpose, we used MCF7 control (MCF7 PLKO) and knockdown of UPR sensors (MCF7 XBP1-KD, MCF7 PERK-KD and MCF7 ATF6-KD) sub-clones of MCF7 (35). We found that BFA-induced increase in the expression of CHOP, pri-miR-616, miR-616-5p and miR-616-3p was attenuated in PERK-knockdown sub-clones of MCF7 (**Fig 4A**), whereas knockdown of ATF6 or XBP1 had no significant effect (**Fig 4B-C**). These results suggest that miR-616 is co-transcribed along with the CHOP transcript and attenuated PERK signalling compromised the induction of CHOP mRNA, pri-miR-616, miR-616-5p and miR-616-3p during UPR.

**Fig 3.**
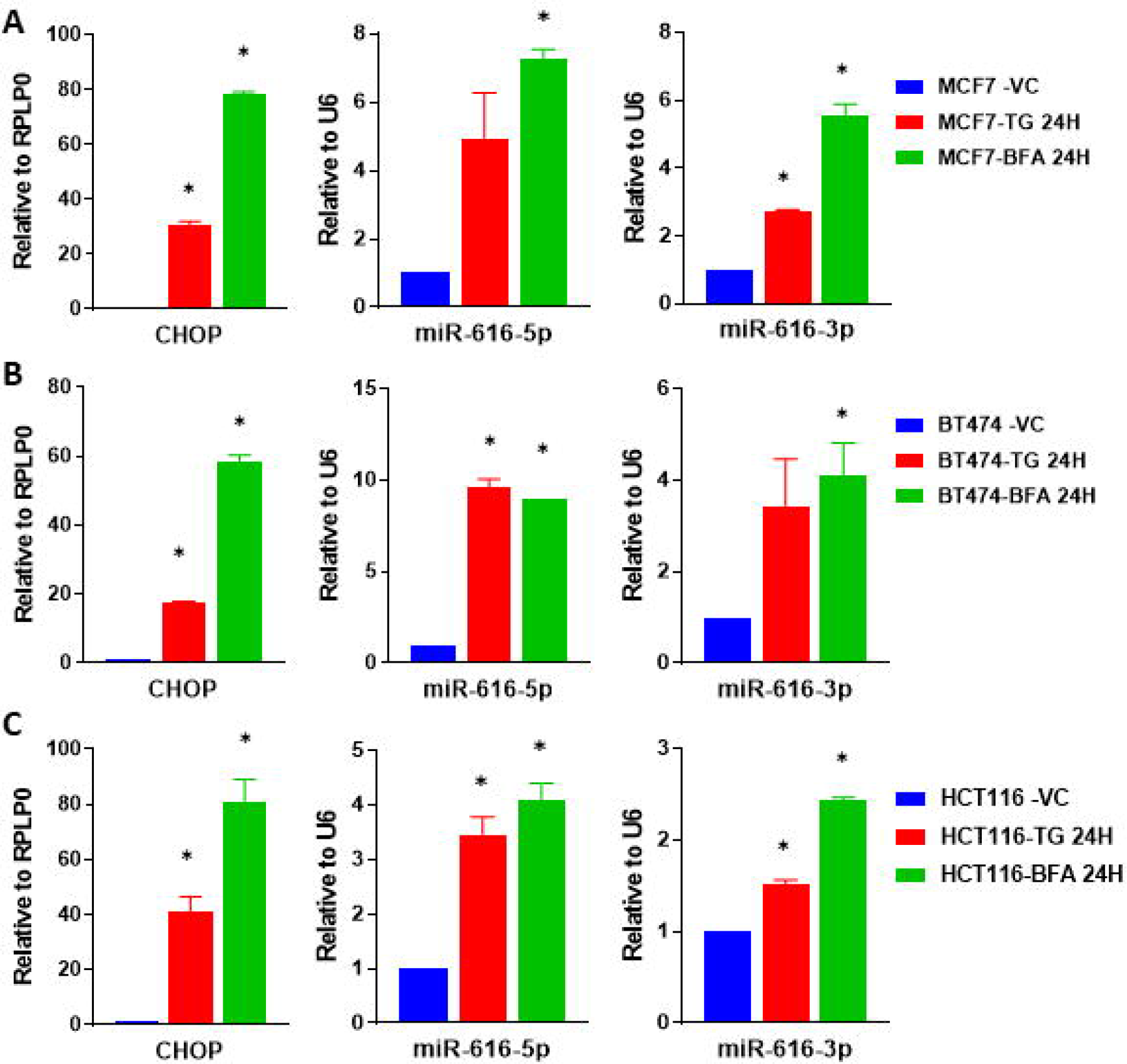
Upregulation of miR-616 (5p and 3p) and its host gene CHOP during UPR. (A) MCF7 cells, (B) BT474 cells and (C) HCT116 cells were treated with (1 μM) TG or (0.5 μg/ml) BFA for 24 hours. Expression of CHOP was quantified by RT-qPCR, normalized using RPLP0. The expression level of miR-616 -5p and -3p was quantified by RT-qPCR, normalized using U6 snRNA. Error bars represent mean ±S.D. from three independent experiments performed in triplicate. *P < 0.05, two-tailed unpaired t-test as compared to untreated control.

**Fig 4.**
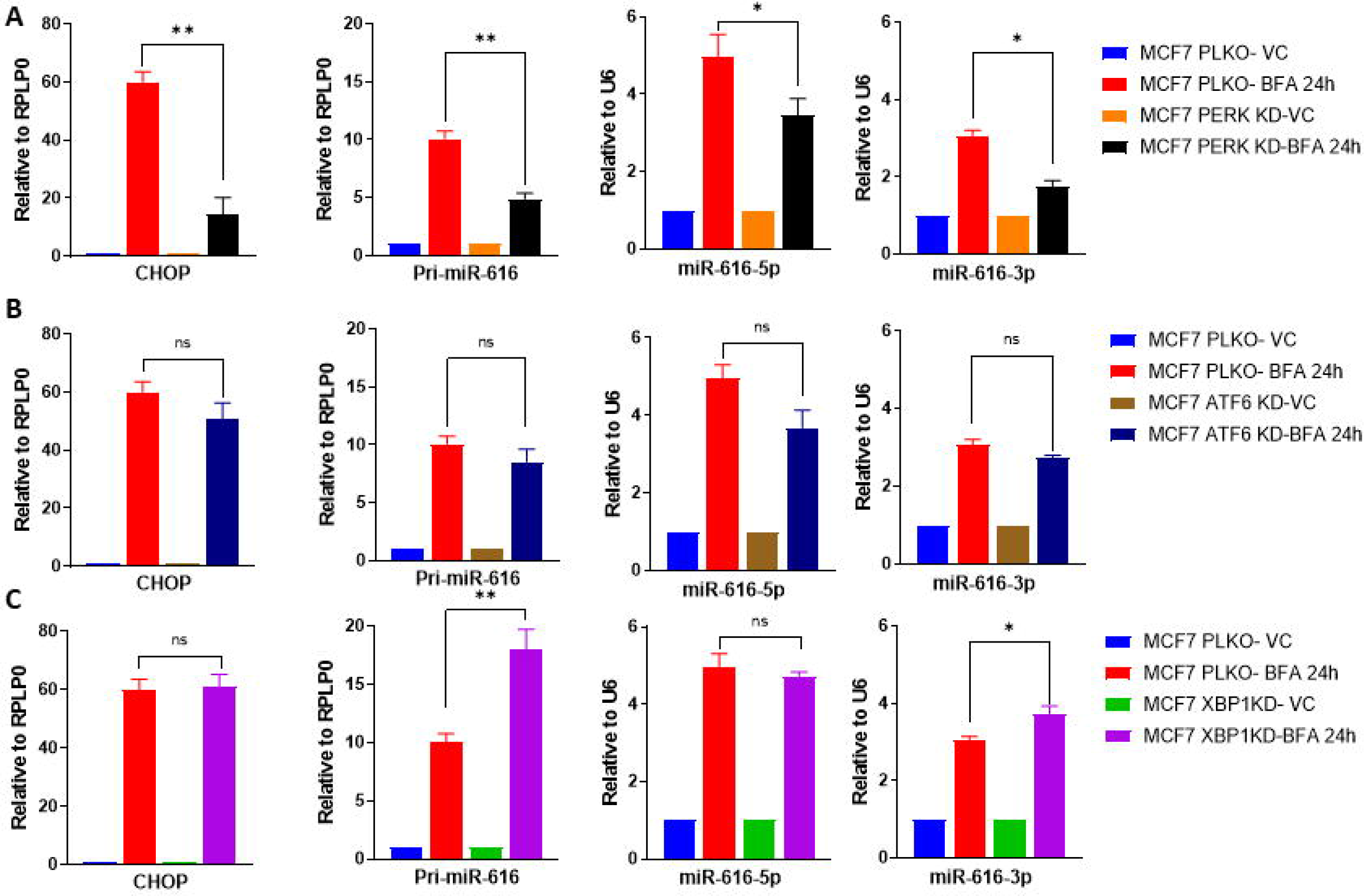
Upregulation of miR-616 during ER stress is dependent on PERK pathway. (A-C) MCF7 PLKO, MCF7 XBP1-KD, MCF7 PERK-KD and MCF7 ATF6-KD sub-clones were treated with (0.5 μg/ml) BFA for 24 hours. Expression level of CHOP and pri-miR-616 was quantified by RT-qPCR, normalizing against RPLP0. The expression level of miR-616 -5p and -3p was quantified by RT-qPCR, normalized using U6 snRNA. *P<0.05, **P < 0.001, two-tailed unpaired t-test. N.S not significant.

### MiR-616 targets c-MYC and reduces cell proliferation of breast cancer cells

Ye et al. reported that ectopic expression of miR-616-3p inhibited growth of MDA-MB231 cells while Arabkari did not find any effect of miR-616 overexpression in MDA-MB231 cells (18) (25). Further, Yuan reported that miR-616-3p promoted growth and migration of MCF7 cells in contrast to Arabkari, who reported a decrease in growth and migration of miR-616 expressing MCF7 cells (19) (25). This could be due to different approaches used for overexpression of miRNAs. Next, we used gain-of-function (overexpression) and loss-of-function (CRISPR-Cas9 KO) approaches in MCF7 cells to study the role of miR-616 in breast cancer. To better mimic the *in vivo* scenario we decided to use a lentiviral system for modulating the expression of both miR-616-5p and -3p concurrently. For this purpose, MCF7 cells were transduced with lentivirus expressing GFP along with miR-616 or Cas-9 along with gRNAs targeting miR-616. The expression of pri-miR-616, miR-616-5p and miR-616-3p was determined in miR-616-overexpressing (MCF7-miR-616-OE) and miR-616-knockout (MCF7-miR-616-KO) sub-clones of MCF7. As expected, the levels of pri-miR-616, miR-616-5p and miR-616-3p were increased in MCF7-miR-616-OE sub-clone (**Fig. 5A**) and decreased in MCF7-miR-616-KO sub-clone (**Fig. 5E**). The MCF7-miR-616-KO sub-clone displayed hypomorphic phenotype with partial reduction in expression of pri-miR-616, miR-616-5p and miR-616-3p. The miR-616 overexpressing (MCF7-miR-616-OE) sub-clone showed significantly decreased cell growth (**Fig. 5B**), while the miR-616 knockout (MCF7-miR-616-KO) sub-clone showed increased cell growth (**Fig. 5F**). This observation was further validated by colony formation assay. The miR-616 overexpressing (MCF7-miR-616-OE) sub-clone of MCF7 cells showed decrease in the number and size of the colonies (**Fig. 5C-D**) but miR-616 knockout (MCF7-miR-616-KO) sub-clone showed an increase in the number and size of the colonies (**Fig. 5G-H**). We found that expression of c-MYC mRNA and protein was reduced in miR-616 overexpressing (MCF7-miR-616-OE) sub-clone of MCF7 cells (**Fig. 6 A and C**) and increased in miR-616 knockout (MCF7-miR-616-KO) sub-clone (**Fig. 6 B and C**). Likewise, overexpression of miR-616 in HCT116 cells significantly decreased cell growth and migration that was accompanied by reduced expression of c-MYC (**S1 Fig**). Our results are in agreement with an earlier report where ectopic expression of miR-616 was shown to inhibit growth of MCF7 cells accompanied by reduced expression of c-MYC (25). Since an inverse relationship between the levels of expression of a miRNA and its target is anticipated, we evaluated expression of c-MYC during conditions of UPR. We found the expression of c-MYC mRNA was decreased in MCF7 (**Fig. 7A**) and BT474 (**Fig. 7B)** cells treated either with TG or BFA. Furthermore, expression of c-MYC protein was decreased in MCF7 and BT474 cells upon treatment with BFA (**Fig. 7C**). These results suggest a role for miR-616 in regulation of c-MYC expression during conditions of UPR.

**Fig 5.**
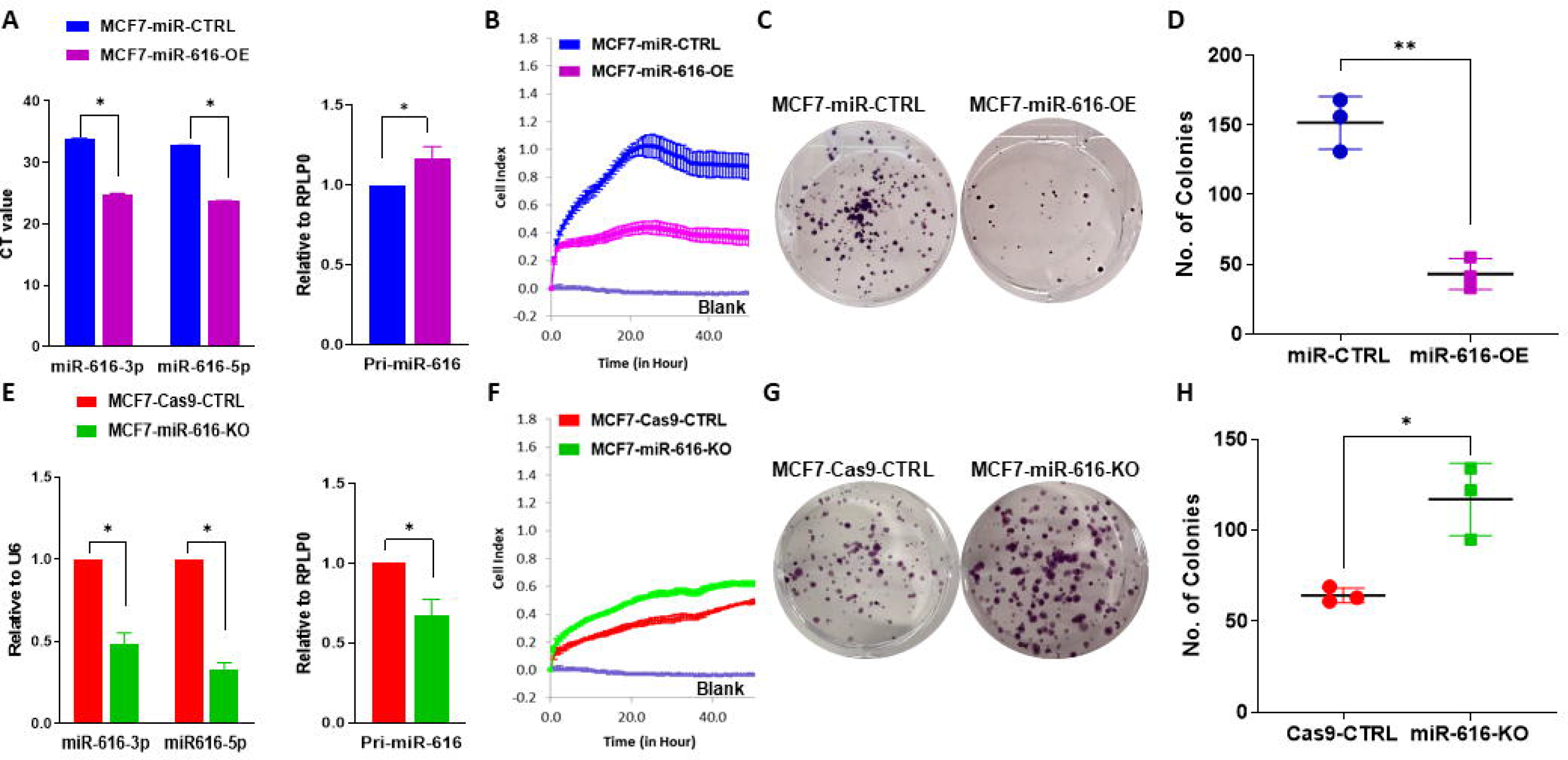
MiR-616 reduces the proliferation of MCF7 cells. (A) The expression of miR-616-3p, miR-616-5p and pri-miR-616 is shown in MCF7-miR-CTRL and MCF7-miR-616-OE sub-clones of MCF7. (B) MCF7-miR-CTRL and MCF7-miR-616-OE cells were plated and cell growth was determined by x-CELLigence. (C-D) MCF7-miR-CTRL and MCF7-miR-616-OE cells were plated in 6-well plate (1000 cells/well) and grown for 14 days. (C) Colonies stained with crystal violet are shown. (D) Quantification of number of colonies for MCF7-miR-CTRL and MCF7-miR-616-OE cells is shown. (E) The expression of miR-616-3p, miR-616-5p and pri-miR-616 is shown in MCF7-Cas9-CTRL and MCF7-miR-616-KO sub-clones of MCF7. (F) MCF7-Cas9-CTRL and MCF7-miR-616-KO cells were plated and cell growth was determined by x-CELLigence. (G-H) MCF7-Cas9-CTRL and MCF7-miR-616-KO cells were plated in 6-well plate (1000 cells/well) and grown for 14 days. (G) Colonies stained with crystal violet are shown. (H) Quantification of number of colonies for MCF7-Cas9-CTRL and MCF7-miR-616-KO cells is shown.

**Fig 6.**
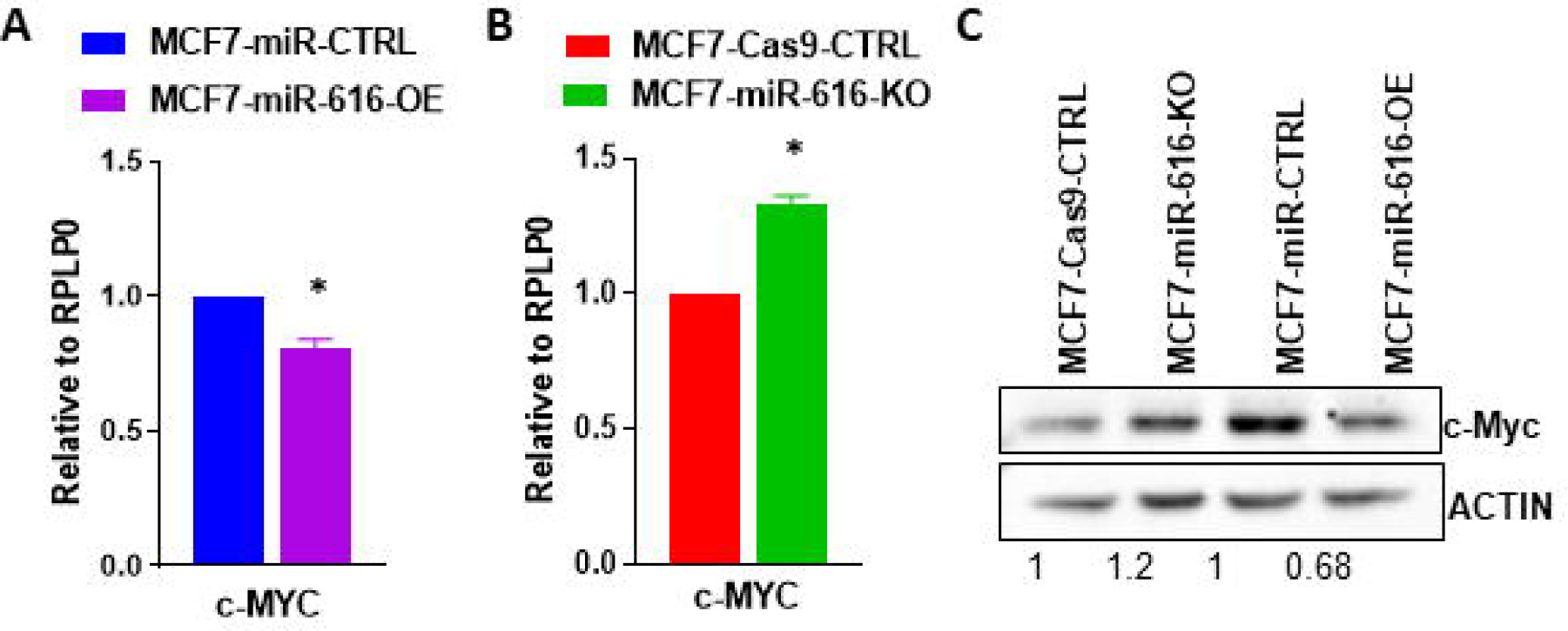
MiR-616 regulates the expression of c-MYC. (A-B) Expression level of c-MYC mRNA is shown in miR-616 knockout and miR-616 overexpressing sub-clones of MCF7 cells as determined by qRT-PCR. Error bars represent mean ± S.D. from two independent experiments performed in triplicate. *P < 0.05, two-tailed unpaired t-test as compared to control. (C) The expression level of c-MYC protein is shown in miR-616 knockout and miR-616 overexpressing sub-clones of MCF7 cells.

**Fig 7.**
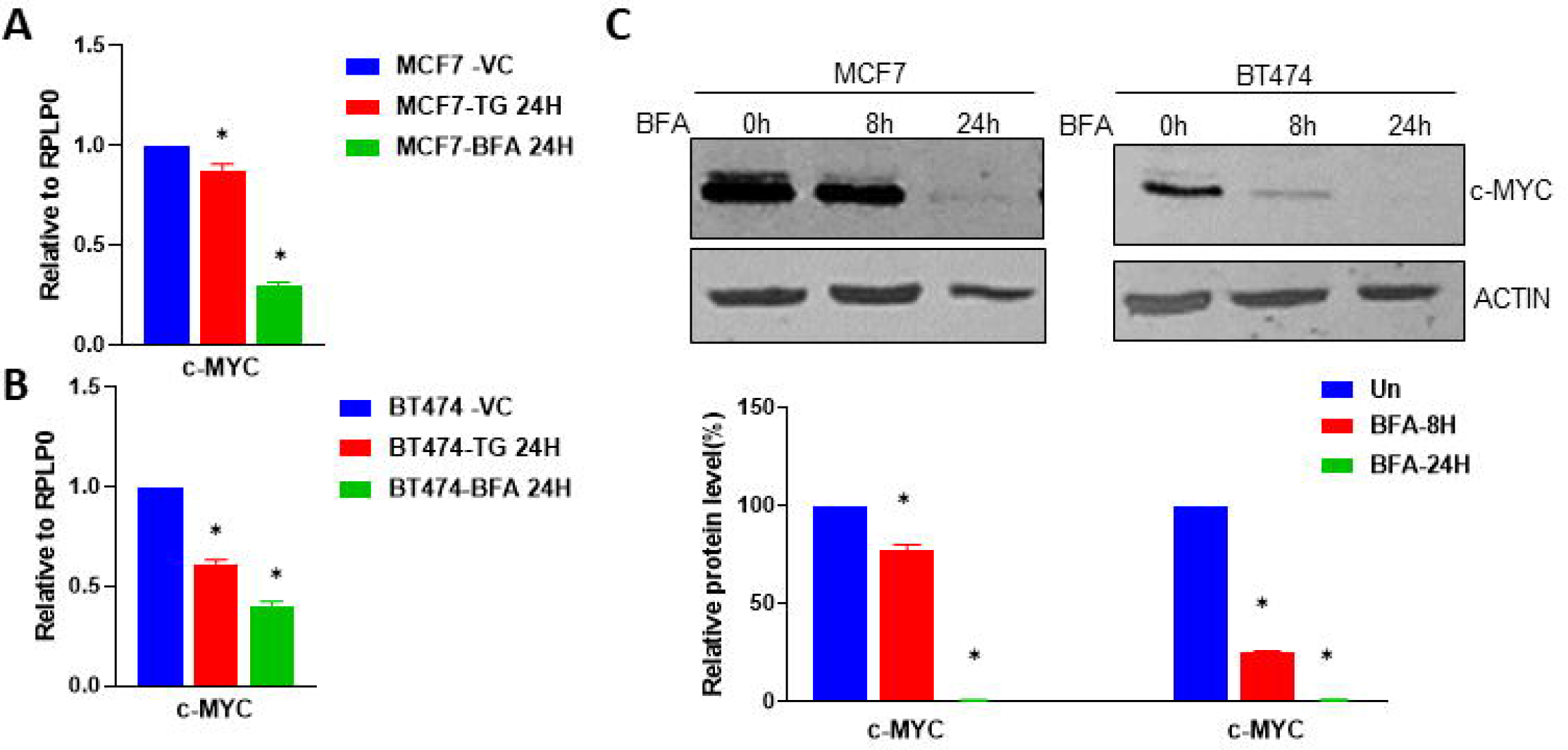
Downregulation of c-MYC during UPR in breast cancer cells. (A) MCF7 cells and (B) BT474 cells were treated with (1 μM) TG or (0.5 μg/ml) BFA for 24 hours. Expression of c-MYC was quantified by RT-qPCR, and normalized using RPLP0. Error bars represent mean ±S.D. from three independent experiments performed in triplicate. (C) MCF7 BT474 cells were treated with (0.5 μg/ml) BFA for indicated time points. Western blotting of total protein was performed using antibodies for c-MYC and actin. Representative Immunoblot and quantification of c-MYC expression normalised to actin are shown (n=3). *P < 0.05, two-tailed unpaired t-test as compared to the untreated control.

### MiR-616 modulates the expression of c-MYC via site in the ORF

To determine the mechanism of c-MYC regulation by miR-616, we used several bioinformatics tools to search for the potential miR-616 binding sites in the reference sequence of c-MYC. TargetScan (36) did not identify miR-616 binding sites in c-MYC. Interestingly, RNAhybrid identified non-canonical miR-616-5p binding sites (at position 77, 2501 and 3076 bp) and miR-616-3p binding sites (at position 614, 917 and 1525 bp) of c-MYC reference Sequence (NM_002467.6) (**S2 Fig**). Furthermore, (miRWALK and RNAhybrid) picked an identical, non-canonical miR-616-3p binding site in the protein-coding region (364-1728 bp) of c-MYC at position 1525 bp (**Fig. 8A-B**). To test whether miR-616 down-regulates c-MYC via this site in coding region, we co-transfected 293T cells with an expression vector containing the ORF of c-MYC and miR-616 plasmid. Cell lysates were collected after 24 h and analysed by immunoblotting. Our results showed that c-MYC protein level was strongly down-regulated by co-expression of miR-616 (**Fig. 8C**). Under similar conditions, co-expression of miR-616 had no effect on protein level of ERα (protein coding region of ERα has no predicted binding sites for miR-616) (**Fig. 8D**). Further, co-expression of miR-4726 (miRNA that lacks binding sites in the protein coding region of c-MYC) and c-MYC had no effect on c-MYC protein level (**Fig. 8E**). Taken together these results suggest that miR-616 regulates the expression of c-MYC most likely via the miR-616 response elements present in its protein coding region.

**Fig 8.**
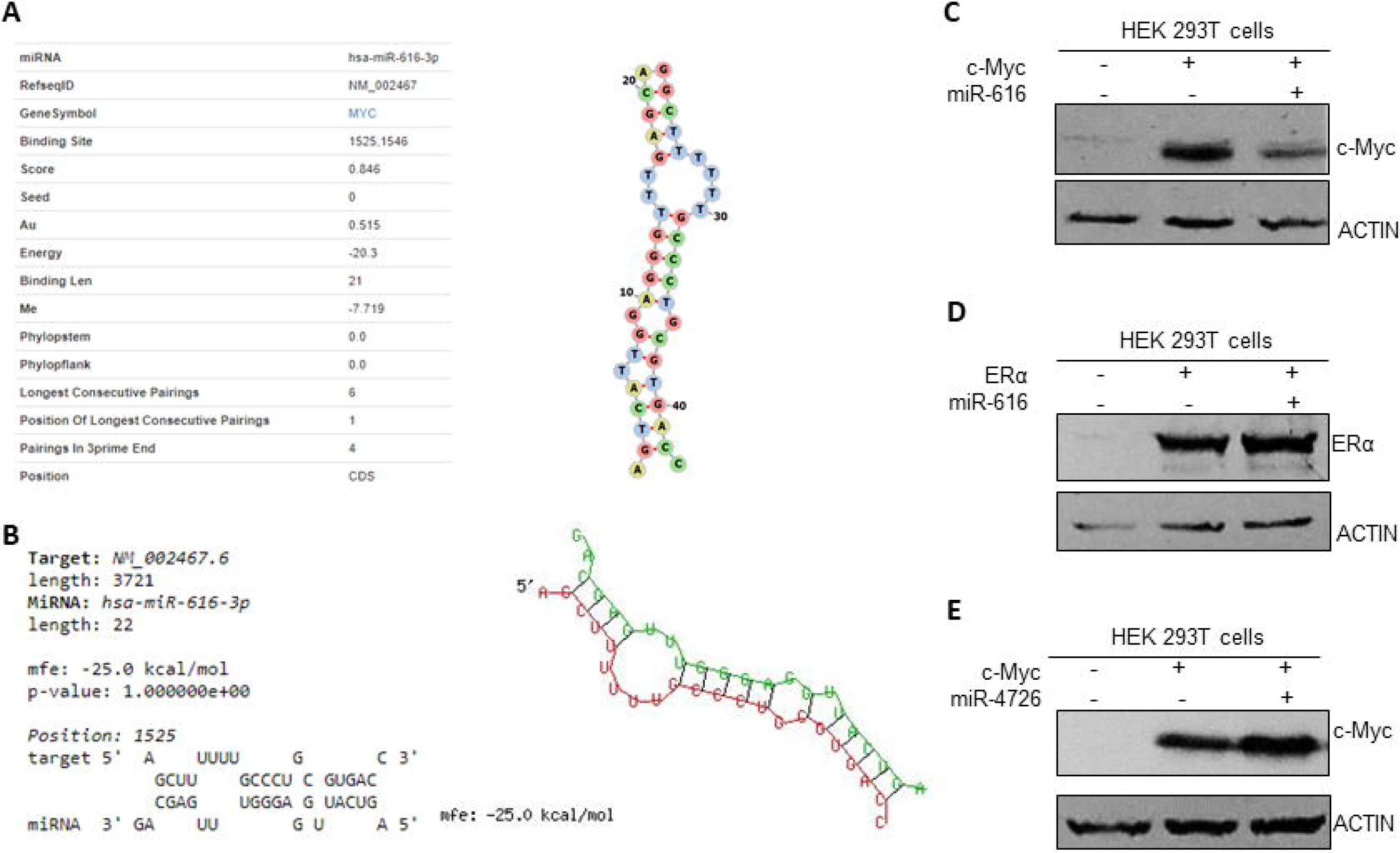
Identification of a miR-616-3p-binding sites in the ORF of c-MYC. Binding site for miR-616-3p identified by (A) miRWALK and (B) RNAhybrid in protein coding region of c-MYC reference sequence is shown. The secondary structure of miRNA-mRNA duplex is shown with miR-616-3p in green and c-MYC mRNA in red. mfe: minimum free energy. (C) 293T cells were transfected with plasmid containing c-MYC ORF in absence and presence of miR-616 plasmid. (D) 293T cells were transfected with plasmid containing ERα ORF in absence and presence of miR-616 plasmid. (E) 293T cells were transfected with plasmid containing c-MYC ORF in absence and presence of miR-4726 plasmid. Western blotting of total protein was performed using the indicated antibodies. 24 h post-transfection western blotting of total protein was performed using the indicated antibodies.

### Expression of c-MYC can reverse the growth inhibitory effects of miR-616

c-Myc is an oncogene frequently overexpressed in human tumours. In light of our results, we reasoned that c-MYC could be the functionally relevant target that mediates tumour-suppressive effects of miR-616. If suppression of c-MYC by miR-616 is indeed crucial for tumour-suppressive effects of miR-616, overexpression of c-MYC should rescue the effect of miR-616 on cell growth. To test this, we transduced lentivirus that express c-MYC into miR-616 expressing MCF7 cells (**Fig. 9A**). We found that expression of c-MYC significantly reversed the growth inhibitory effects of miR-616 in MCF7 cells (**Fig. 9B**). Further, we observed that expression of c-MYC rescued the inhibitory effects of miR-616 on the number and size of the colonies in MCF7 cells (**Fig. 9C**). Next we determined whether c-MYC can rescue the effects of miR-616 on migration of cells. The bright field images of the wound healing revealed that expression of c-MYC restored the rate of wound healing in miR-616 expressing MCF7 cells that was comparable to control cells (**Fig. 9D-E**). Thus overexpression of c-MYC rescued the inhibitory effects of miR-616 on cell growth, colony formation and migration. The data therefore suggests an essential role for c-MYC as a mediator of the biological effects of miR-616 in breast cancer cells.

**Figure 9.**
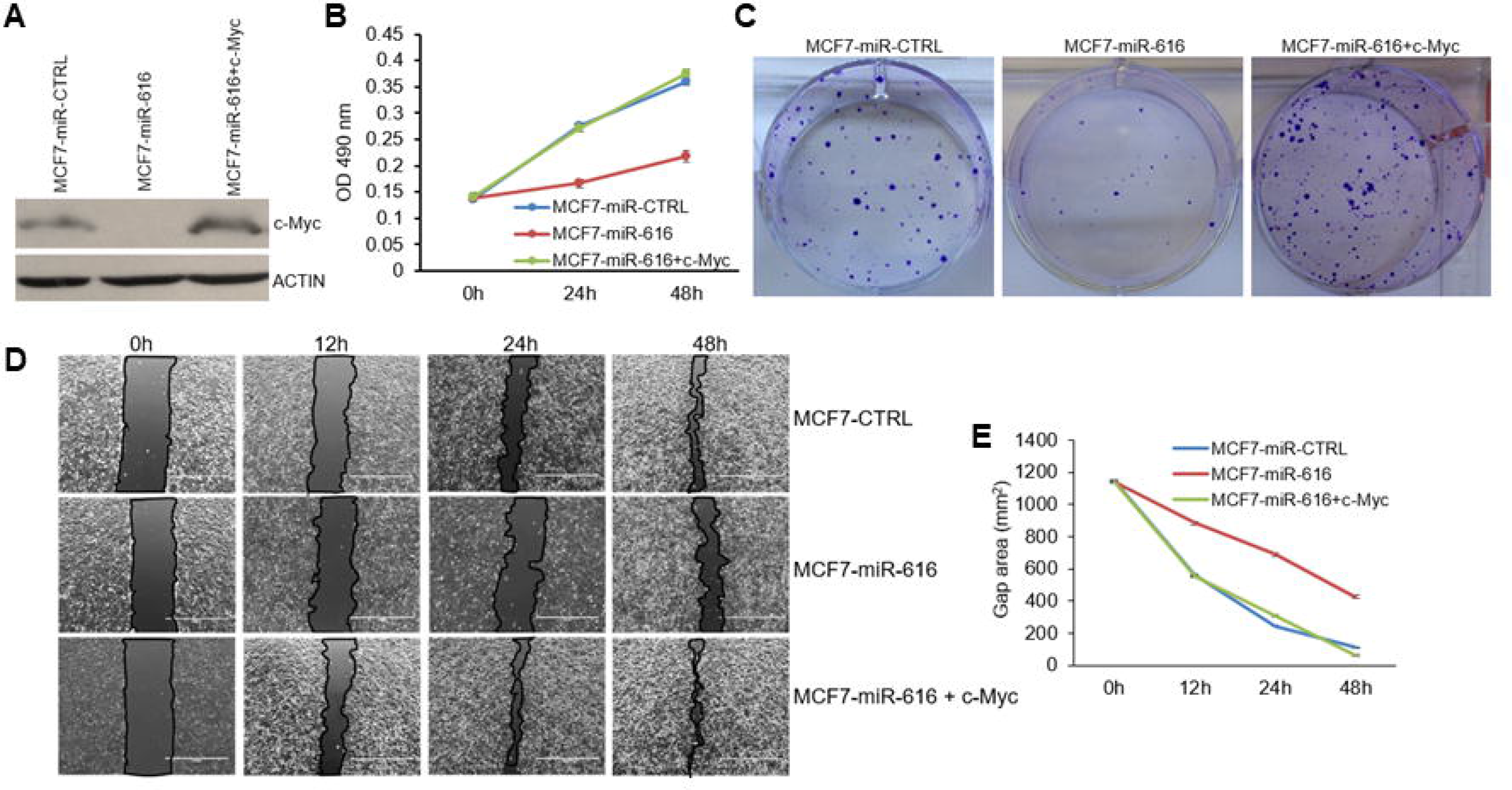
Restoration of c-MYC expression overcomes the inhibitory effect of miR-616 on cell proliferation. (A) The expression level of c-MYC protein in MCF7-miR-CTRL and MCF7-miR-616 and MCF7-miR-616 cells transduced with c-MYC lentivirus is shown. (B-C) Cell growth was determined by MTS assay and colony formation assay using MCF7-miR-CTRL and MCF7-miR-616 and MCF7-miR-616 cells expressing c-MYC. (B) Cell proliferation was assessed by MTS assay. Line graphs show the absorbance in cells at the indicated time points. Error bars represent mean ± S.D. from three independent experiments performed in triplicate. (C) Cells were plated in 6-well plate (1000 cells/well) and grown for 14 days. Colonies stained with crystal violet are shown. (D-E) Cells were plated in 6-well plate. The monolayer of cells was scratched with a micropipette tip (200 μl). Black lines indicate the wound borders at indicated time points post-scratching. Scale bar, 1000 μM. Line graph shows the scratch gap quantified using Image J software at the indicated time points. Error bars represent mean ±S.D. from three independent experiments. *P < 0.05.

### Prognostic value of the miR-616 in human cancers

We assessed the prognostic value of miR-616 in different cancer types using KM plotter. We found that expression of miR-616 was associated with longer overall survival (OS) in cervical squamous cell carcinoma, Head-neck squamous cell carcinoma, uterine corpus endometrial carcinoma and Thymoma (**S3 Fig**). Surprisingly, the increased expression of miR-616 was associated with shorter OS in esophageal adenocarcinoma, sarcoma, renal clear cell carcinoma, lung squamous cell carcinoma and hepatocellular carcinoma (**S3 Fig**). Next, we used TNMplot to determine the expression of CHOP and miR-616 in normal and tumour tissue samples of breast. We found that expression of CHOP and miR-616 was reduced in tumour samples as compared to tumour adjacent normal tissue (**Fig. 10A-B**). Further, the expression of miR-616 was associated with poor OS in luminal A subtype and better OS HER2 subtype of breast cancer (**Fig. 10C**).

**Fig 10.**
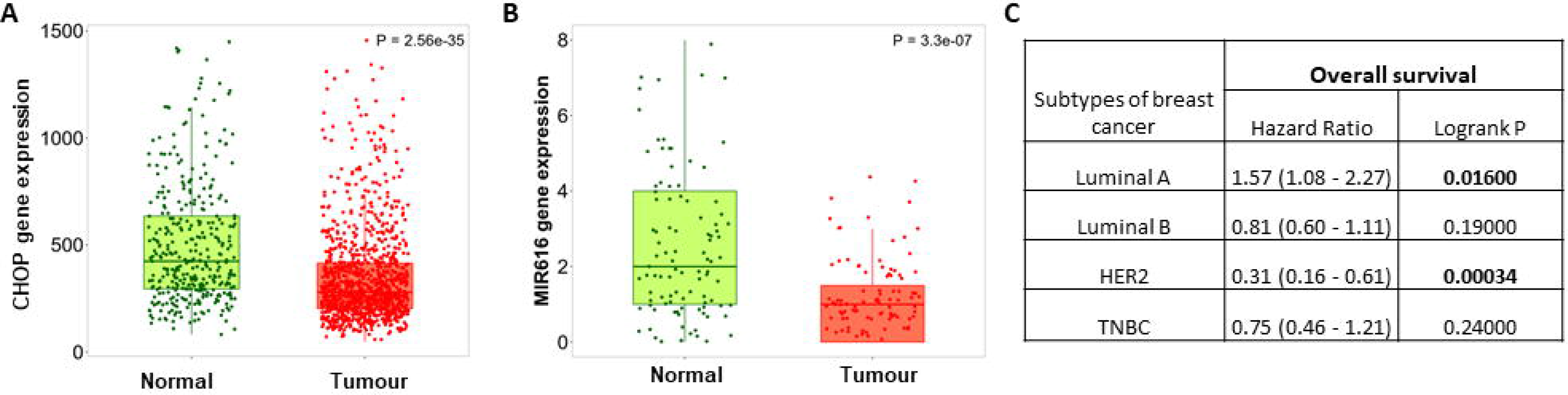
Prognostic value of miR-616 in subtypes of breast cancer. TNMplot (https://tnmplot.com/analysis/) was used to determine the expression of CHOP and miR-616 in human breast cancer. (A) Box plot for expression of CHOP in tumour (n=1097) and normal (n=403) tissues for human breast cancers is shown. (B) Box plot for expression of miR-616 in paired tumour (n=112) and tumour adjacent normal (n=112) for human breast cancers is shown. (C) KM Plotter (https://kmplot.com/) was used to determine the association of miR-616 with overall survival (OS) in subtypes of breast cancer.

## DISCUSSION

MiRNAs regulate the wide range of cellular processes because of their ability to alter post-transcriptional gene expression (37). We have evaluated the role of UPR regulated miRNA-616 in cell growth and proliferation. We have shown that expression of miR-616 and its host gene (CHOP) is upregulated during ER stress in a PERK-dependent fashion (**Fig 4**). ER stress and UPR activation contributes to the initiation and progression of human diseases, such as cancer, neurodegenerative and cardiovascular disease (2, 38). We observed concordant expression of the CHOP and the mature miR-616 which suggests their transcriptional co-regulation (**Fig 3, 4**). Our results suggest a mechanism for increased expression of miR-616 in patho-physiological conditions where role of UPR is implicated.

Several studies have reported that miR-616 acts as a tumour promoting miRNA in human cancers (11, 31, 39). However, our results show that miR-616 expression attenuates the proliferation of MCF7 cells while reduced expression of miR-616 increases it (**Fig 5**). Furthermore, we observed that expression of miR-616 was reduced in human breast cancers as compared to tumour adjacent normal tissue (**Fig 10**). In agreement with our results miR-616-3p has been shown to potentiate apoptosis in HUVECs (16), inhibit head and neck squamous cell carcinoma progression (17) and inhibit cell growth and mammosphere formation of breast cancer cells (18, 25). Any small RNA having G-rich 6mer seed sequence (nt 2-7 of the guide strand) can reduce viability of cells when associated with RNA-induced silencing complex. G-rich seed sequence mediates toxicity by targeting C-rich seed matches in the 31 UTR of genes critical for cell survival referred to as Death Induced by Survival gene Elimination (DISE) (30). A systematic study to evaluate DISE activity of all 4096 possible 6mer seed sequences in six cell lines (three human and three mouse) revealed the mechanism underlying this toxicity and a web-based algorithm (https://6merdb.org) can predict the activity of an RNA with a known 6mer seed (40). Analysis of seed sequences from both 5p and 3p strands of miR-616 revealed that both strands can exhibit opposing effects on viability in a cell-type dependent manner (**S4 Fig**). Interestingly, several miRNAs belonging to miR-371-373 Cluster (hsa-miR-371a-5p, hsa-miR-371b-5p, hsa-miR-372-5p and hsa-miR-373-5p) that share seed sequence with miR-616-5p, have been shown to reduce the progression and metastasis of colon cancer (41) as well as induction of cell cycle arrest (42). Taken together, these observations indicate the presence of context-dependent oncogenic and tumour suppressive effects of miR-616 in human cancers. Indeed context-dependent tumour-promoting and tumour-suppressive role has been documented for several miRNAs such as miR-17-92 cluster (43, 44). This cluster maps to a region frequently amplified in Burkitt’s lymphoma, diffuse large B-cell lymphoma, follicular lymphoma and lung cancer (26, 43). The expression of miR-17-92 is reduced in prostate cancer (17) and miR-17-92 expression in intestinal epithelial cells inhibited colon cancer progression by suppressing tumour angiogenesis (17).

Studies on target recognition by miRNAs have shown that sequence complementarity at the 5’ end of the miRNA, the so-called “seed region” at positions 2 to 7, is a main determinant for target recognition (36). However, a perfect seed match of its own is not a good predictor for miRNA regulation and a number of studies have shown that miRNA can regulate the expression of target genes via the sites with a G: U wobble and/or mismatch in the seed region (36). Further, a study using a cross-linking and immunoprecipitation method to experimentally identify microRNA target sites in an unbiased manner has reported a significant number of non-canonical sites (45, 46). Hence, presence of perfect seed complementarity is not essential for regulation of target genes by miRNA. In addition, target sites for endogenous miRNAs have been reported in ORFs and 5⍰ UTRs, but they are less frequent than those in the 3’ UTR (47, 48). We found a non-canonical miR-616-3p binding site in the protein coding region of c-MYC transcript (**Fig 8**). Several studies have reported the presence of miRNA-binding sites in the protein coding sequence of the genes. Indeed miRNAs have been shown to regulate embryonic stem cell differentiation, DNA methylation, regulation of apoptosis, aortic development, and tumour suppression via the functional miRNA-binding sites in the protein coding sequence of the target genes (47, 48, 49). Several studies have reported the upregulation of c-MYC in different cancer cells and tumours such as neuroblastomas, lung, and breast cancer (50). The expression of c-MYC was downregulated in miR-616 expressing and upregulated in miR-616 knockout MCF7 cells (**Fig 6**). These observations suggest that effects of miR-616 on cell growth are primarily mediated by downregulation of c-MYC.

PERK has been shown to act as a haploinsufficient tumour suppressor, where the nature of its function is determined by gene dose (51, 52). Transient pause in protein synthesis due to eIF2α phosphorylation by PERK is beneficial by reducing secretory load in the ER. Phosphorylation of NRF2 by PERK attenuates Keap1-mediated degradation of NRF2 and promotes expression of anti-oxidant enzymes through the antioxidant response elements (53, 54). PERK signalling can upregulate the CHOP/DDIT3 transcription factor, which inhibits expression of the gene encoding anti-apoptotic BCL-2 to hasten cell death, in addition to enhancing the expression of pro-apoptotic BCL-2 members such as BIM (54). Studies in both cellular and animal models with CHOP gene deficiency have shed light on the pro-apoptotic role of CHOP during cellular stress (54). Further, ectopic expression of CHOP was reported to induce a G1 cell cycle arrest (55). CHOP maintains the integrity of the human hematopoietic stem cell (HSC) pool by eliminating HSCs harbouring oncogenic mutations, and decrease the risk of leukaemia. CHOP induction triggers apoptosis of premalignant cells to prevent malignant progression in a mouse lung cancer model. Hepatocyte-specific CHOP ablation increased tumourigenesis in high fat diet-induced steatohepatitis and Hepatocellular carcinoma (56, 57). CHOP has been shown to promote cancer progression, when fused with FUS/TLS or EWS protein by genomic rearrangement (58, 59). The FUS-CHOP oncoprotein has been shown to induce metastasis in in vivo model of sarcoma (58). Accumulating data suggest that CHOP impinges upon several aspects of cancer including initiation as well as progression of tumours (54). Our results show that miR-616 supressed cell growth of cancer cells through suppressing c-MYC expression. Our results establish a new and unexpected role for the CHOP locus by providing evidence that miR-616 can inhibit cell proliferation by targeting c-MYC.

## ACKNOWLEDGEMENTS

We are grateful to the Technical Officers and administrative team in Pathology, School of Medicine at University of Galway, Ireland.

## Supporting information

S1 Fig. **miR-616 reduces the proliferation of HCT116 cells**. (A) Control (HCT116-miR-CTRL) and miR-616-overexpressing (HCT116-miR-616) sub-clones of HCT116 cells were plated and cell growth was determined by MTS assay. Line graphs show the absorbance in cells at the indicated time points. Error bars represent mean ± S.D. from three independent experiments performed in triplicate. (B) HCT116-miR-CTRL and HCT116-miR-616 cells were plated in 6-well plate (1000 cells/well) and grown for 14 days. Colonies stained with crystal violet are shown. (C) HCT116-miR-CTRL and HCT116-miR-616 cells were plated in 6-well plate. The monolayer of cells was scratched with a micropipette tip (200 μl). Black lines indicate the wound borders at indicated time points post-scratching. Scale bar, 1000 μM. Line graph shows the scratch gap quantified using Image J software at the indicated time points. (D) The expression level of c-Myc protein is shown in HCT116-miR-CTRL and HCT116-miR-616 cells. *P < 0.05, two-tailed unpaired t-test as compared to the control sub-clone.

S2Fig. **Identification of a miR-616-5p and miR-616-3p -binding sites in the c-Myc**. RNAhybrid analysis of c-MYC reference sequence and miR-616. The secondary structure of miRNA-mRNA duplex is shown with miRNA in green and mRNA in red. mfe: minimum free energy.

S3 Fig. **Association between miR-616 expression and overall survival in human cancer**. Web-based algorithm KM plotter (http://kmplot.com/analysis/) was used to evaluate the association between miR-616 with overall (OS) survival.

S4 Fig. **Prediction of the effect of miR-616-5p and miR-616-3p on cell viability through their 6mer seed sequence**. The activity of 6-mer seed of was evaluated in three human (HeyA8-ovarian, H460-lung and H4-brain) and three mouse (M565-liver, 3LL-lung and GL261-brain) cell lines. MiRNAs with low viability (high toxicity) are shown in red and with high viability (low toxicity) are shown in green. Predominantly expressed mature miRNAs (guide miRNA) are shown in dark purple while lesser expressed (passenger miRNA) are shown in light purple. The miRNAs sharing seed sequence with miR-616-5p are also shown.

