## Supplementary figures and images for "PERK-mediated induction of miR-5p and miR-3p arms of miR-616 regulates cell growth by targeting c-MYC"

### Supplementary Figure 1

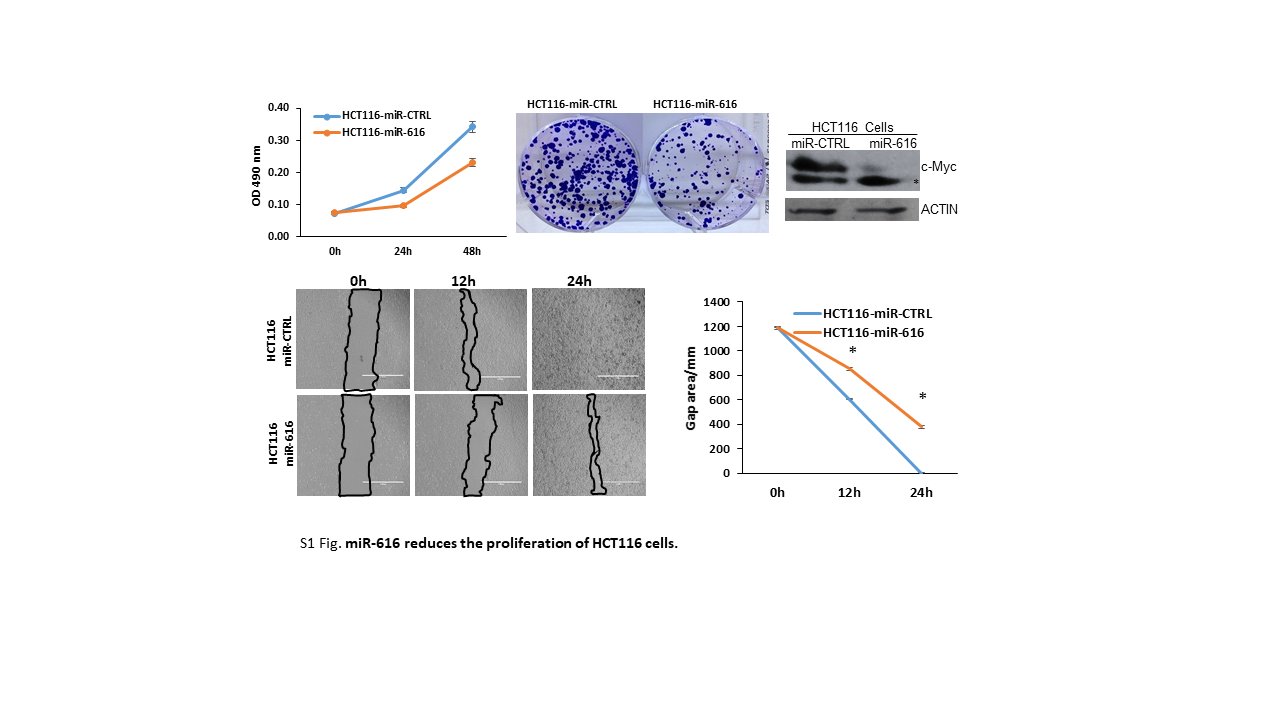

### Supplementary Figure 2

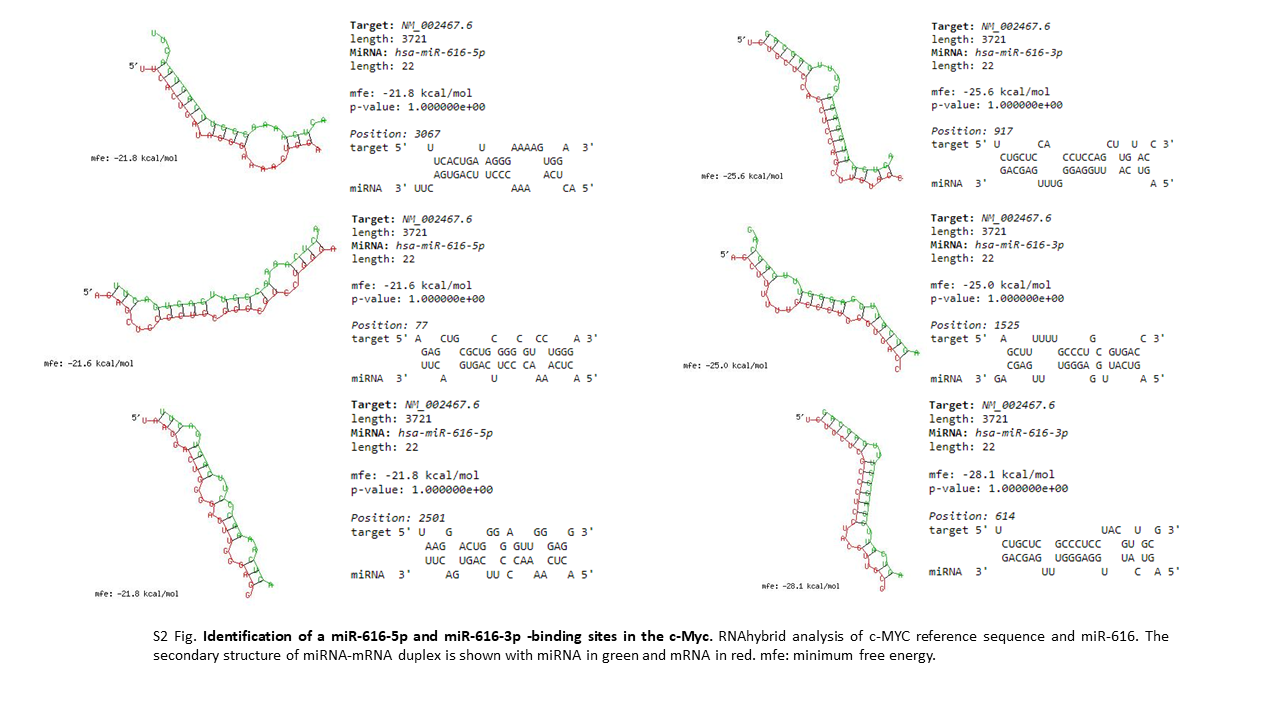

### Supplementary Figure 3

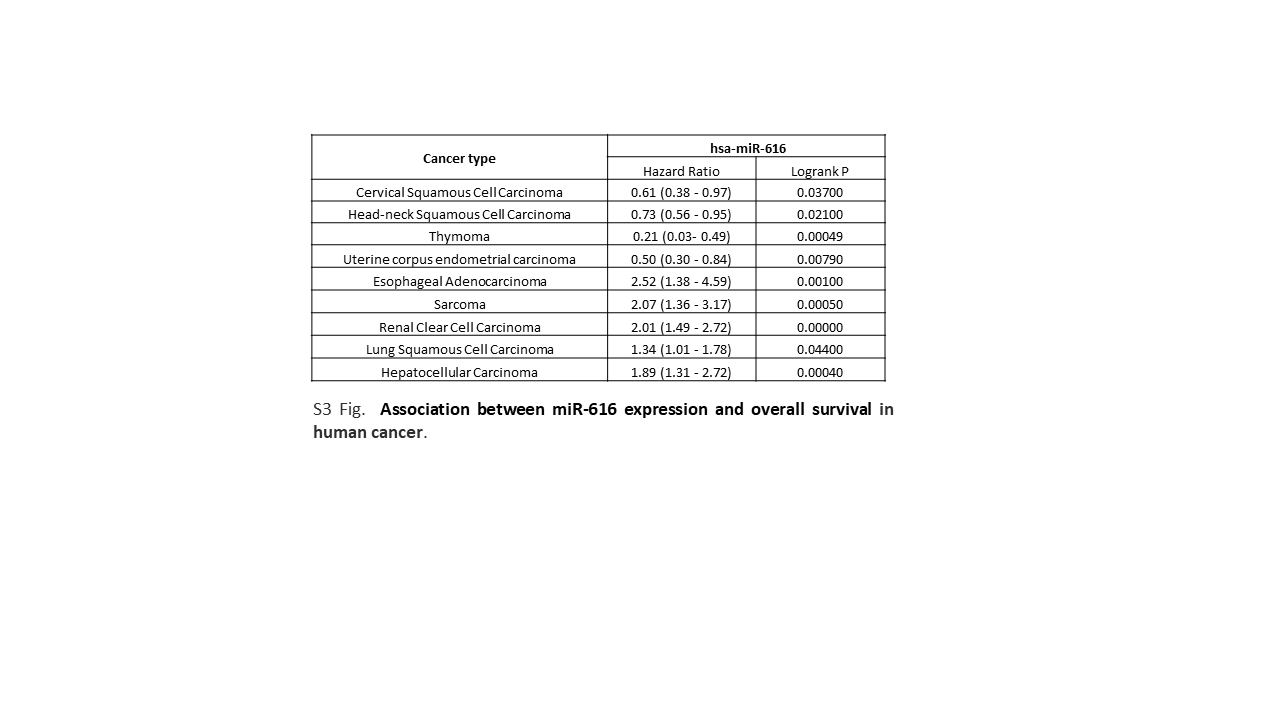

### Supplementary Figure 4

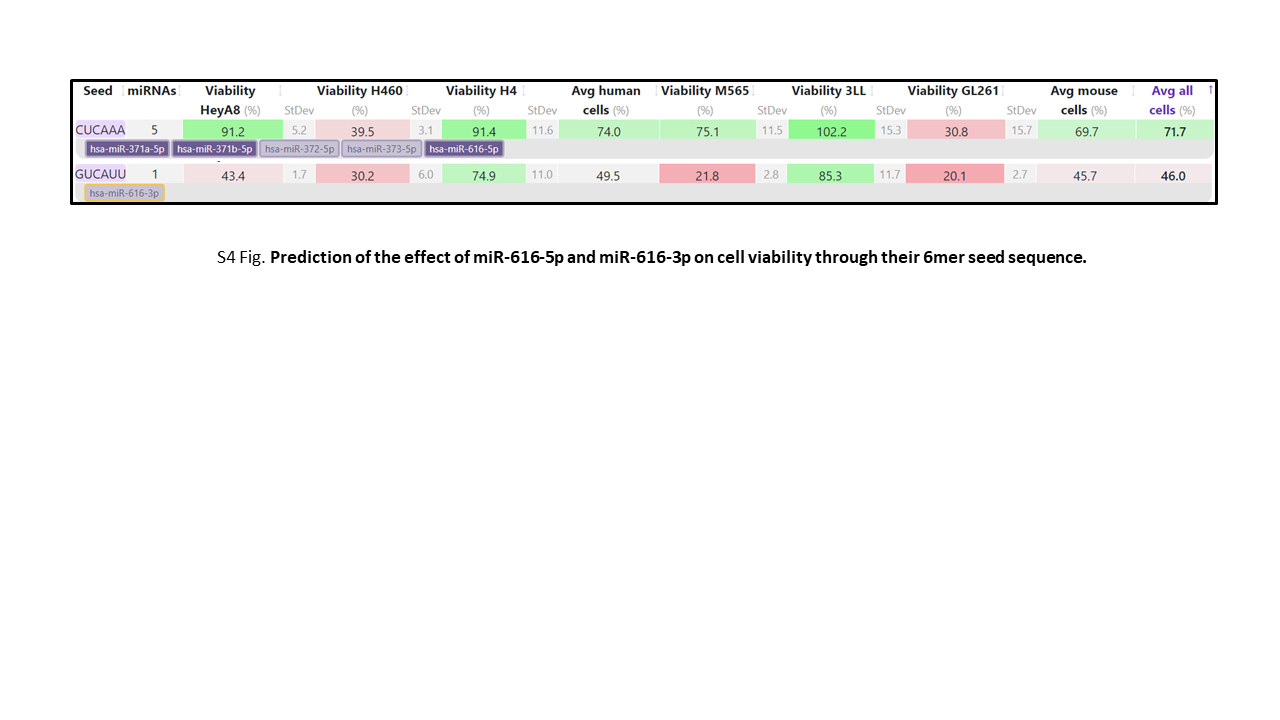
